# PEPSI: Polarity measurements from spatial proteomics imaging suggest immune cell engagement

**DOI:** 10.1101/2023.10.13.562299

**Authors:** Eric Wu, Zhenqin Wu, Aaron T. Mayer, Alexandro E. Trevino, James Zou

**Affiliations:** Department of Electrical Engineering, Stanford University, Stanford, CA, USA; Enable Medicine, Inc., Menlo Park, CA, USA; Department of Biomedical Data Science, Stanford University, Stanford, CA, USA

**Keywords:** subcellular localization, proteomics, multi-plex immunofluorescence

## Abstract

Subcellular protein localization is important for understanding functional states of cells, but measuring and quantifying this information can be difficult and typically requires high-resolution microscopy. In this work, we develop a metric to define surface protein polarity from immunofluorescence (IF) imaging data and use it to identify distinct immune cell states within tumor microenvironments. We apply this metric to characterize over two million cells across 600 patient samples and find that cells identified as having polar expression exhibit characteristics relating to tumor-immune cell engagement. Additionally, we show that incorporating these polarity-defined cell subtypes improves the performance of deep learning models trained to predict patient survival outcomes. This method provides a first look at using subcellular protein expression patterns to phenotype immune cell functional states with applications to precision medicine.

## 1. Introduction

Spatial proteomics methods such as immunofluorescence (IF) and immunohistochemistry (IHC) enable an unprecedented view of tumor microenvironments by preserving the spatial structure of tissues at subcellular resolution^1^. However, standard analyses aggregate and average protein expression within segmented single cells, discarding sub-cellular and morphological signals^2^. This approach introduces a number of analytical limitations. First, segmentation can be imprecise. Second, subcellular protein expression patterns could allow the inference of cellular functional states (i.e. polarized vs. uniform). Thus, while cells can be phenotyped in the context of their spatial neighbors, cells that exhibit differential protein localization are not differentiated.

The relationship between protein localization and function is well-established in many contexts. For instance, during T cell engagement with presented antigens (e.g., on tumor cells), the CD4 and CD8 coreceptors are recruited to the immune synapse, while they present uniformly on the surface of a cell in a naive or exhausted state ^3–5^. The immune synapse, also known as the supramolecular activation cluster, is a specialized junction formed by many proteins during an immune response. In the context of a tumor immune response, active and engaged T cells could correlate with better patient survival, whereas the presence of exhausted or inactive T cells may be indicative of worse outcomes ^6–8^. However, it is unknown to what extent this or other such dynamic subcellular localization events are discernable from whole slide scale histology images.

Common analyses of cell morphology ^9–13^ utilize computer vision models for the automatic extraction of image features from tissue patches. However, interrogating such models for specific cell-cell interactions is difficult. Previous work toward characterizing surface protein localization includes statistical methods for identifying ligand-receptor pairs in transcriptomics ^14,15^, polarity localization measurements in mRNA ^16,17^, and co-localization with protein expression ^18^.

In this paper, we present a novel approach, PEPSI (**P**rotein **E**xpression **P**olarity **S**ubtyping in **I**mmunostains), for measuring subcellular protein localization toward characterizing the tumor microenvironment. We describe a simple, explainable method for computing the polarity of cell surface biomarkers. We apply this metric on multiple large-scale CODEX (co-detection by indexing) datasets spanning over two million cells, three clinical sites, and 600 patient samples.

We focus on several key immune cells that are well-characterized and known to express polarized surface protein markers during activation/engagement. We define additional cell subtypes relating to morphology (polarized, uniform) for representative biomarkers (CD8, CD4, CD20) of immune cells (T cells and B cells). We find that surface protein marker polarity is significantly correlated with positive patient outcomes, even after controlling for various technical artifacts, suggesting that this may be important for characterizing the functional state of immune cells. We believe that inferring functional subtypes of cells can offer a better understanding of patient response to drug treatments and disease prognostic indicators.

## 2. Results

### 2.1. Polarity measurement

We describe a straightforward method for extracting polarity measurements for a given cell based on a polar transformation of the IF signal with respect to the cell centroid (**Figure 1A, Methods**). Plotting the distribution of scores for four markers in their cognate expression cell types - CD8 in CD8 T cells, CD4 in CD4 T cells, CD20 in B cells, and PanCK in tumor cells - shows that the scores exhibit continuous distributions (**Figure 1B**). The first three biomarkers, which are known to polarize in cells undergoing immune activation, show significantly higher average polarity scores versus PanCK, which is not known to polarize as such. To obtain a discrete polarity classification, we threshold the raw scores based on an empirical heuristic (**Methods**), obtaining proportions for polar, uniform, and ‘other’ cells for each of the three immune cell types (**Figure 1C**). For instance, polar cells account for 3.7%, 3.0%, and 2.6% of CD8 T cells, CD4 T cells, and B cells, respectively. Example cells were randomly inspected in their cell contexts to visually validate the classifications (**Figure 1D**).

**Fig. 1:**
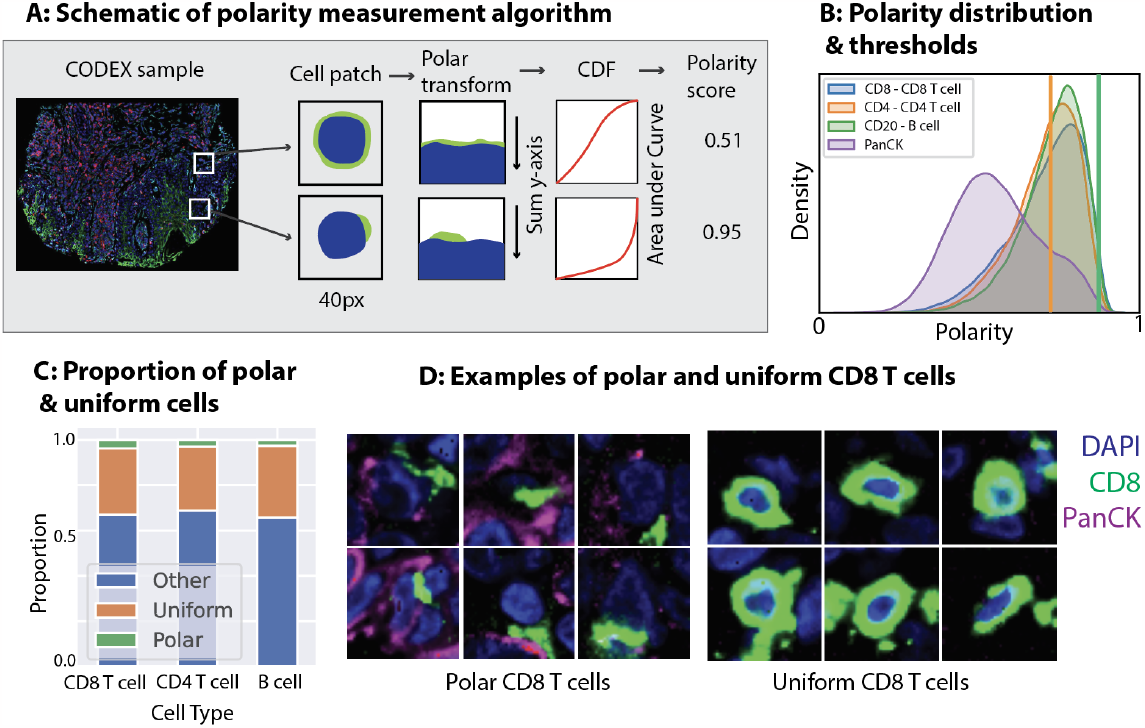
Overview of the PEPSI polarity measurement framework. **Panel A:** Schematic of polarity measurement algorithm. For a given mIF sample, patches (40px by 40px) are extracted around each cell. For each cell, a polar transform is computed on the patch, followed by summing along the y-axis and then computing the area under the CDF curve, yielding a polarity score (from 0 to 1). **Panel B:** The polarity score histograms are shown for CD8, CD4, CD20, and PanCK biomarkers in CD8 T cells, CD4 T cells, B cells, and tumors, respectively. The orange (left) line indicates the upper threshold chosen for identifying uniform cells, whereas the green (right) line indicates the lower threshold chosen for identifying polar cells. Cells in between two thresholds are indicated as ‘Other’. **Panel C:** The percent proportions of polar, uniform, and other cells across the three cell types and their key biomarkers. **Panel D:** For CD8 T cells, representative examples of polar and uniform CD8 T cells are shown, and color-coded by relevant biomarkers.

### 2.2. Polarized cell neighborhoods are more enriched with tumors

Next, we explore whether polar immune cells might exhibit differences in their cellular neighborhoods with respect to uniform cells of the same type. Examples of a polarized CD8 T cell adjacent to a tumor cell (left) or not (right) are shown in **Figure 2A**. We found that, for CD8 T cells, CD4 T cells, and B cells, tumor cells were consistently enriched in the immediate cell neighborhoods of polar cells versus uniform cells (**Figure 2B**). Conversely, we also find that cells with tumor cell neighbors are more likely to be polar (**Supp. Table 1)**. Given that polar expression can indicate antigen engagement during contact with tumor cells ^19^, this provides evidence that polarity is a biologically significant biomarker.

**Table 1:** Adding polarity-specific cell types improves patient survival prediction in machine learning models. To validate the usefulness of the polarity-specific cell types derived from our polarity measurement method, we train two models to predict patient survival status, with and without the additional cell types (polar and uniform CD8 T cells, CD4 T cells, B cells). Each cell represents the AUC of the model’s prediction of patient survival. We observe that adding the additional cell types improves model performance across three datasets, three clinical sites, and two disease types. Standard deviations of bootstrapped samples are reported in parentheses. Predictions are generated at the sample level.

| Survival status | UPMC-HNC | Stanford-CRC | DFCI-HNC (using UPMC-HNC model) generalization |
| --- | --- | --- | --- |
| MLP |  |  |  |
| - Baseline | 0.759 (0.053) | 0.538 (0.132) | 0.655 (0.074) |
| - Adding polarity-defined cell subtypes | **0.803 (0.049)** | **0.559 (0.135)** | **0.672 (0.072)** |
| GNN |  |  |  |
| - Baseline | 0.839 (0.048) | 0.684 (0.092) | 0.853 (0.046) |
| - Adding polarity-defined cell subtypes | **0.856 (0.045)** | **0.743 (0.090)** | **0.880 (0.051)** |

**Fig 2:**
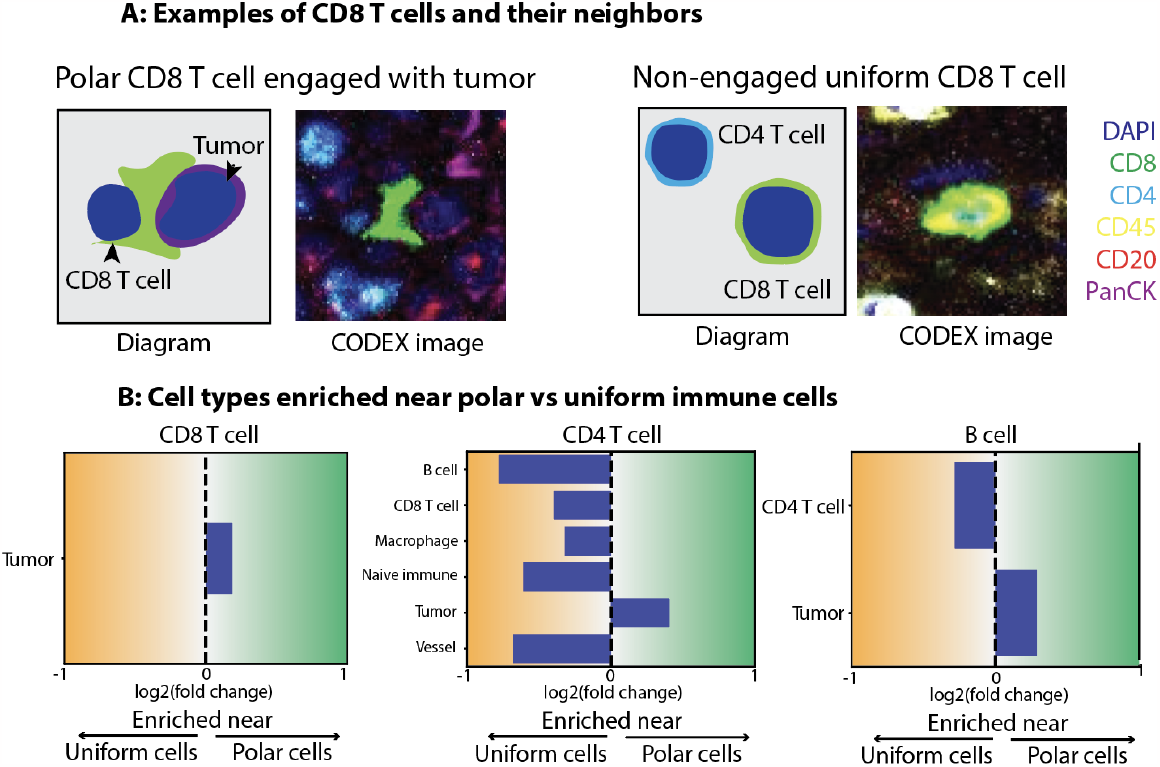
Tumor cells are more present next to polar cells versus uniform cells. **Panel A:** Diagrams and mIF images illustrating two possible states of CD8 T cells. Left: A CD8 T cell engaged with a tumor cell, with polar expression of CD8 at the immunological synapse. Right: A uniformly expressed CD8 T cell, with no tumor engagement. **Panel B:** Since polarity may be indicative of tumor engagement, we measure the cell type composition of neighborhoods around polar versus uniform cell types. We find that tumor cells are consistently and significantly more enriched in polar cell neighborhoods versus uniform cell neighborhoods for all three immune cell types. We compute bootstrapped 95% confidence intervals for each neighboring cell type and only show cell types with significant log fold changes.

### 2.3. Metric control experiments

In addition to visual validation, we perform tests to validate that the metric distribution is not explained by simple technical covariates or noise. We find that polarity cannot be simply explained by significantly more crowded cell neighborhoods (**Supp Figure 1A**) or differences in cell size (R2 of 0.08, **Supp Figure 1B**). During antigen engagement, multiple biomarkers are known to jointly express at the site of the immunological synapse ^20^. **Supp. Figure 1C** measures the correlation of polarity scores between all pairs of biomarkers as expressed in all T cells, B cells, and tumor cells. CD3e, a biomarker known to express during engagement, is jointly polarized with CD4 and CD8, while, PanCK, a biomarker not known to be active during antigen engagement, does not correlate with CD20, CD3e, CD4, or CD8. We note that the observed co-polarity between CD4 and CD8 is likely due to expression from neighboring cells that are being captured by our algorithm as originating from the same cell, an artifact that occurs in a small fraction of T cells (**Supp. Figure 1D)**.

### 2.4. Polarity-defined cell types improve model prediction of survival outcomes

To demonstrate that the newly classified polar or uniform cell subtypes have biological or clinical relevance, we utilize deep learning models to predict patient survival from cell phenotypes. We train two models: a 3-layer multi-layer perceptron (MLP) neural network, which takes as input the percent composition of cell types per sample and predicts a binary outcome (five-year survival); and a graph-based neural network (GNN) that takes as input a 3-hop neighborhood of cells centered around a single cell, and predicts the survival status of the sample from which the neighborhood of cells originated. Both models show modest but consistent improvement in performance across three distinct studies and two disease types after including the 6 new cell types (**Table 1**). **Supp. Table 2** shows ablations where the MLP model is trained on each polar/uniform cell type individually. Of note, a model is trained with Ki67 polarity in tumor cells as a negative control (since Ki67 is not known to express polarly) and demonstrates no improvement over the baseline. Finally, we use the percent of polar cells per sample and compute the AUROC in its usefulness in predicting survival outcomes in **Supp. Table 3** and find that even this simple metric alone has predictive accuracy above chance.

### 2.5. Presence of polar cells improves patient survival with in silico models under label and spatial permutations

In the permutation experiments shown in **Figure 3** and **Supp. Table 4**, the GNN model predicts significantly worse survival in tumor microenvironments where the subtype of the immune cells are flipped from polar to uniform **(Figure 3B)**. The inverse is also true; the predicted survival improves when cells are flipped from uniform to polar. Even when fixing the cell type composition, dispersing the location of the immune cells away from the tumor cells results in a decrease in predicted survival (and vice versa) **(Figure 3C)**. These results suggest that polar immune cells are important not simply for their presence in a sample, but for their proximity to tumor cells.

**Fig 3:**
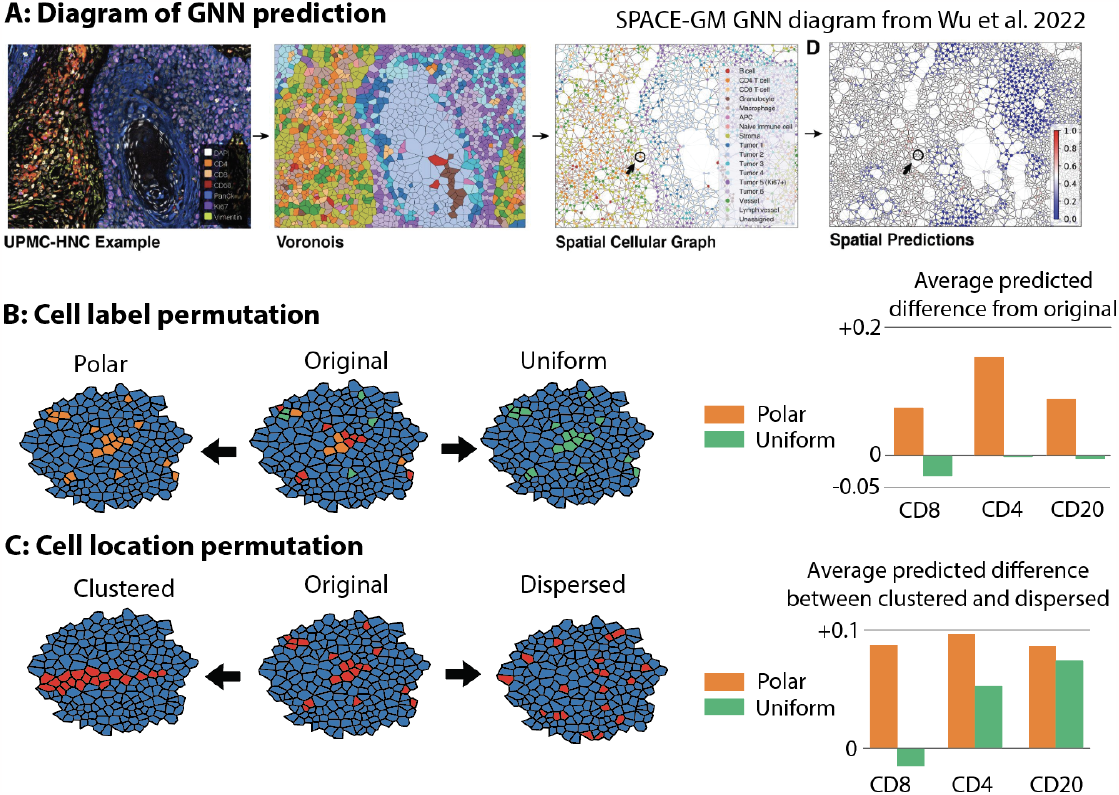
*In silico* experimentation reveals that polar cells are correlated with positive patient outcomes. **Panel A:** A schematic of a graph neural network described in *Wu et al. 2022*. A mIF sample is represented by a Voronoi diagram, which is projected into a spatial graph. A graph neural network is trained to predict survival outcomes based on 3-hop cellular neighborhoods. **Panel B:** Using a trained GNN, we perform label permutation on each sample graph, where the subtype of each immune cell is flipped to either polar or uniform, and the averaged model prediction is measured. Even when fixing spatial neighborhoods, we observe an increased predicted survival probability when cells are polarized, and a slight decrease when cells are turned into uniform states. **Panel C**: Now, fixing the cell types, polar and uniform cell neighborhoods are sampled and spatially permuted. We observe, on average, a larger increase in predicted survival when polar cells are dispersed from the clustered state than with uniform cells.

## 3. Methods

### 3.1. Datasets

Our primary dataset consists of 308 samples from 81 patients with head and neck squamous cell carcinomas at the University of Pittsburgh Medical Center (UPMC-HNC). Two external validation datasets are used: a colorectal cancer dataset with 292 samples from 161 patients from Stanford University (Stanford-CRC) to demonstrate generalization to another disease; and a head and neck squamous cell carcinomas dataset with 112 samples from 29 patients from Dana Farber Cancer Institute (DFCI-HNC) to demonstrate generalization to an additional clinical site. The number of samples, patients, coverslips, and total cells in each dataset is described in **Supp. Table 5A**. Phenotype annotations for UPMC-HNC are described in **Supp. Table 5B**. Full CODEX data acquisition and preparation details are described in **Supp. Methods**. UPMC-HNC is chosen as the primary training and evaluation dataset as it contains the largest number of samples, coverslips, and total cells. We evaluate our models on held-out coverslips not seen during training to assess model robustness to technical artifacts across coverslips.

The UPMC-HNC and Stanford-CRC datasets have one held-out coverslip for model validation and one held-out coverslip for model evaluation. The Stanford-CRC dataset has half of one coverslip randomly split and held out for model validation and one held out for model evaluation. The DFCI-HNC dataset has one coverslip randomly split by patients for model evaluation.

### 3.2. Biomarker expression preprocessing

Single-cell expression was computed for each biomarker by 1. applying a deep learning cell segmentation algorithm (DeepCell) ^21^ on the DAPI biomarker channel (nuclear stain) to obtain nuclear segmentation masks; 2. successively dilating segmentation masks by flipping pixels each time with a probability equal to the fraction of positive neighboring pixels (repeated 9 times); 3. computing the mean expression value across pixels within the single cell; and 4. normalizing the expression values across all cells in a sample using quantile normalization and arcsinh transformation followed by a z-score normalization:

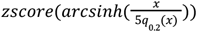

Where *zscore* is defined given μ and σ, the mean and standard deviation across all cell expression values in the sample:

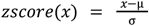

*x* is the vector of a biomarker’s values in a sample, *arcsinh* is the inverse hyperbolic sine function; and *q*0.2(*x*) is the 20th percentile of *x*.

### 3.3. Image patch generation

After preprocessing (tile & cycle alignment, stitching, deconvolution, and background correction) CODEX data is available as multichannel OME-TIFF files, with each image channel corresponding to the fluorescence signal (expression) of a distinct biomarker probe. To prepare the input image patches for the deep learning model, we perform the following: All pixel values for a biomarker in a sample are normalized using ImageJ’s AutoAdjust function.

### 3.4. Cell type ground truth and predictions

To produce cell type labels, we first obtained a cells-by-features biomarker expression matrix - for each marker, we took the average signal across all pixels in a segmented cell. This matrix was normalized and scaled as described above, and then principal component (PC) analysis was performed. We constructed a nearest-neighbor graph (k = 30) of cell expression in PC space with the top 20 PCs, then performed self-supervised graph clustering ^22^ on the result. Clusters were manually annotated according to their cell biomarker expression patterns. This procedure was performed on a subset of 10,000 cells and subsequently used to train a kNN algorithm. This algorithm was used to transfer labels to the entire dataset. The cell type labels that were used are: Tumor (CD15+, CD20+, CD21+, Ki67+, Podo+, Other), Naive immune cell, Granulocyte, Vessel, CD4 T cell, Macrophage, CD8 T cell, Stromal / Fibroblast, APC, Lymph vessel, and B cell.

### 3.5. Calculating polarity score

Our polarity measurement methodology is described in **Figure 1**. The segmentations are used to calculate cell center coordinates. For each cell, a 40px square patch is extracted around the center pixel. Several de-noising steps are first taken: 1. low/background values are zeroed out (values < 0.1), 2. biomarker expressions that spatially overlap with the DAPI channel are subtracted out in both the center and neighborhood cells.

We then transform the patch from cartesian coordinates to polar coordinates using the scikit package (skimage.transform.warp_polar). The polar image is then summed along the y-axis, producing a 1-dimensional vector. An additional refining step is taken where cells are assigned ‘other’ if the vector 1. sums to 0 either along the x-or y-axis, 2. does not contain multiple unique values, or 3. has a mean less than 0.02. Finally, the vector is normalized within a [0,1] range and sorted in ascending order, and a score is computed by subtracting the AUC of the sorted vector from 1.

On its own, the polarity score is difficult to interpret and incorporate into existing analysis pipelines that rely on discrete cell phenotypes. Thus, we define three cell subtypes based on the polarity score value: uniform (cells with a polarity score below a threshold), polar (cells with a polarity score above a threshold), and other (cells that fall in between both thresholds). To obtain optimal thresholds for defining polar and uniform cell types from the polarity scores, we perform a two-dimensional grid search on the MLP model and select the pair of values that yielded the highest validation AUC score in the survival prediction task. From this process, we obtain 0.94 and 0.8 as the polar and uniform threshold cutoffs, respectively (**Figure 1B). Figure 1C** shows the polarity distributions after thresholding.

These thresholds are used to define six new cell subtypes for polar and uniform CD8 T cells, CD4 T cells, and B cells. CD8, CD4, and CD20 were used as the representative surface biomarkers for each of the three cell types, respectively. These three cells and biomarkers were chosen as they are known in the literature to exhibit polar expression during engagement ^19^.

### 3.6. Machine learning models

We train two machine learning models to evaluate the benefit of including the six newly defined cell subtypes. First, we use a 3-layer multilayer perceptron (MLP) neural network that accepts the percent composition of cell types per sample and predicts binary 60-month survival. Each layer contains 256 nodes followed by a LeakyReLU ^23^ activation function. Each model is trained with binary cross-entropy loss across 200 epochs and a learning rate of 0.001.

Second, we train a graph-based neural network (GNN) ^24^ that takes as input 3-hop cell neighborhoods and predicts neighborhood-level survival status (**Figure 2A)**. This model transforms the structure of each sample into a graph network, where cells are connected by edges to neighboring cells. It then pools information about the neighboring cells’ cell types to output an outcome probability score for each cell. The sample predictions are generated by averaging the scores across all cells in that sample. Model training details follow the procedures described in Wu et al. ^24^.

Each of the models is trained first on the original 16 cell types (baseline) and then trained using the 6 additional cell types. In both of these settings, each model is trained and evaluated on the UPMC-HNC and Stanford-CRC datasets. An additional evaluation is performed on the DFCI-HNC dataset using the UPMC-HNC trained model.

### 3.7. Permutation experiments

To assess the effect of polar/uniform cell types on the GNN model’s survival predictions, we perform several permutation experiments. In the first experiment (**Figure 3A)**, we flip the cell type label of all immune cells to either polar or uniform and evaluate the predicted survival probability in each scenario. In the second experiment, we sample random subgraphs containing immune cells and tumor cells for CD4 T cells, CD8 T cells, and B cells. Then, we flip all immune cells to either polar or uniform and perform a spatial permutation, where we shuffle the immune cells into either clustered (where all immune cells are neighbors) or dispersed (where immune cells are randomly located in the subgraph) orientations and evaluate the predicted probabilities for each orientation.

## 4. Discussion

We describe a robust, interpretable subcellular morphology metric that reflects macro-biological states. Although our results do not conclude that these polarity events definitively quantify immune synapses, they do suggest that such measurements represent biologically relevant signals in tumor microenvironments. Though our described method is performed on CODEX data, it can similarly be applied to other lower-plexed imaging techniques like IHC that include a nuclear marker (i.e. DAPI) and one or more surface biomarkers.

To date, there has not been prior consensus demonstrating that biological events like engagement and activation or exhaustion can be reliably observed at the standard resolution of mIF imaging. Potential confounders include bleedover, sample slicing artifacts, measurement noise, cell size, and density of neighboring cells. We address this by conducting several negative control experiments and find that these factors alone do not adequately explain the signal present in our polarity measurements.

One counter-hypothesis is that polarity measurements serve as a proxy for neighborhood information -- i.e. the presence of certain cell types or spatial arrangements. Another possibility is that they are primarily an imaging artifact (for instance, irregular borders due to slicing). To test these, we trained a GNN that incorporates local neighborhood information into its predictions and then introduced the polar and uniform cell types. The results show that the new cell types improve performance even in models that have access to cell neighborhood information, suggesting that it introduces information beyond the neighborhood cell type composition or spatial arrangement of cells.

Further experimental evidence is required to define these observations as a specific biological phenomenon, i.e. T cell engagement. However, we believe that this work provides evidence of the importance of measuring and incorporating subcellular polarity information into tissue microenvironment analyses, and represents an important step toward a personalized understanding of disease states, drug response, and patient prognosis.

## Supplementary Materials

All supplementary tables, figures, and data are available at: https://docs.google.com/document/d/1n97PEC2kq41fNOWMXrOASyd42DZ0_nUapnEs6HoeJ8c

## Code Availability

Code for replicating the experiments in this paper is present in this code repository: https://gitlab.com/enable-medicine-public/polarity

