## Supplemental Tables and Figures for "PEPSI: Polarity measurements from spatial proteomics imaging suggest immune cell engagement"

#### Supplementary Tables and Figures:

**Supplementary Table 1:** CD8 T cells with at least one immediate neighbor that is a tumor cell are significantly more likely to be polar cells and significantly less likely to be uniform cells. Bootstrapped 95% confidence intervals are reported in parentheses.

| CD8 T cells: | % of polar cells | % of uniform cells |
|---|---|---|
| **Cells with tumor neighbor** | 4.2% (4.1%, 4.4%) | 28.8%, (28.5%, 29.1%) |
| **Cells without tumor neighbor** | 3.5% (3.4%, 3.7%) | 32.6%, (32.3%, 32.9%) |

**Supplementary Figure 1:** Negative control experiments on potential confounding variables for polarity measurements. In these experiments, we verify that polarity is not simply a proxy for cell statistics, like the number of neighboring cells (**Panel A**; black bars represent standard deviation**)** or cell size (**Panel B)**. We also verify that the polarity measurements are jointly positively correlated in key biomarkers that we expect to co-express (e.g. CD3e/CD8), and not correlated with biomarkers we do not expect to co-express (i.e. PanCK) (**Panel C)**. We confirm that the high correlation between CD4 and CD8 polarity is likely due to signal bleed over of one of the biomarkers from neighboring cells **(Panel D)**.

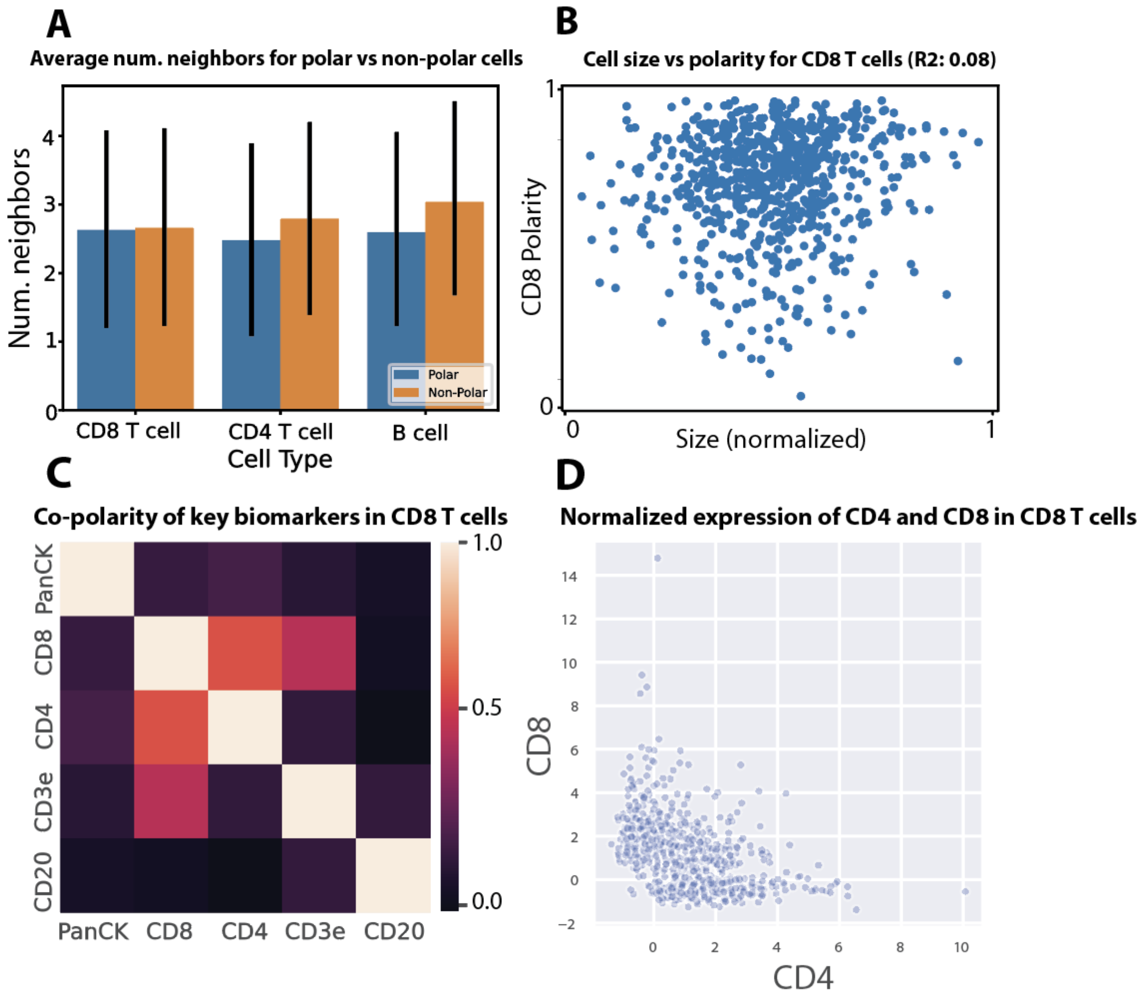


**Supplemental Table 2:** MLP model ablations. In addition to **Table 1**, we test the effect of adding polar and uniform cell subtypes for each cell type individually. For instance, "Adding CD8" represents training a model with the addition of two new cell subtypes (polar and uniform CD8 T cells). Each cell reports the AUC of the model's prediction on the UPMC-HNC test dataset. Of note, each of the three immune biomarkers (CD8, CD4, CD20) improves the model above the baseline, but Ki67 (a tumor biomarker) does not.

| **Survival status**<br>3-layer MLP | **UPMC-HNC** |
|---|---|
| - Baseline | 0.767 |
| - Adding CD8 | 0.766 |
| - Adding CD4 | 0.770 |

| | |
|---|---|
| - Adding CD20 | 0.797 |
| - Adding Ki67 | 0.752 |
| - Adding CD8, CD4, and CD20 | 0.805 |

**Supplementary Table 3:** AUC of polarity vs survival outcome. To test whether polarity alone provides a predictive signal for patient survival, we compute the AUC using the polarity score as a prediction probability. We find that most combinations of biomarkers and cell types have significant but weak predictive power. Bootstrapped 95% confidence intervals are reported in parentheses.

| Cell Type | Biomarker | AUC (95% CI) |
|---|---|---|
| CD8 T cell | CD4 | 0.624 (0.557, 0.687) |
| CD8 T cell | CD8 | 0.594 (0.522, 0.663) |
| CD8 T cell | CD3e | 0.560 (0.497, 0.627) |
| CD4 T cell | CD4 | 0.586 (0.522, 0.651) |
| CD4 T cell | CD8 | 0.545 (0.477, 0.613) |
| CD4 T cell | CD3e | 0.572 (0.512, 0.635) |

**Supplementary Table 4:** Full results from the permutation experiments described in **Figure 3**. Each cell contains the average predicted probability of survival across all test subgraphs. Rows labeled *Change* indicate the difference from the original prediction (unpermuted).

| | CD8 T cell | CD4 T cell | B cell |
|---|---|---|---|
| **Cell Label Permutations** | | | |
| - Original | 0.45 | 0.45 | 0.45 |
| - Polar | 0.52 | 0.60 | 0.54 |
| - *Change* | +0.07 | +0.15 | +0.09 |
| - Uniform | 0.42 | 0.45 | 0.44 |
| - *Change* | -0.03 | 0.00 | 0.01 |
| **Cell Location Permutations** | | | |
| ***- Polar cell neighborhood*** | | | |
| - Clustered | 0.72 | 0.80 | 0.68 |

| | | | |
|---|---|---|---|
| - Dispersed | 0.80 | 0.86 | 0.74 |
| - *Change* | +0.08 | +0.06 | +0.06 |
| - ***Uniform cell neighborhood*** | | | |
| - Clustered | 0.71 | 0.77 | 0.63 |
| - Dispersed | 0.70 | 0.79 | 0.67 |
| - *Change* | -0.01 | +0.02 | +0.04 |

**Supplemental Table 5:** Dataset descriptions of the three datasets analyzed.

**A: Descriptions of the three CODEX datasets.**

| Dataset | No. of samples | No. of patients | No. of coverslips | No. of total cells |
|---|---|---|---|---|
| UPMC-HNC | 308 | 81 | 8 | 2,061,102 |
| Stanford-CRC | 292 | 161 | 4 | 632,280 |
| DFCI-HNC | 112 | 29 | 1 | 259,620 |

**B: Phenotype annotations for UPMC-HNC dataset.**

| Dataset | $N_{sample}$ | $N_{patient}$ | $N_{batch}$ | $N_{cell}$ | Phenotype Annotations | | | | |
|---|---|---|---|---|---|---|---|---|---|
| | | | | | Primary Outcome | Survival Length | Recurrence | Recurrence Interval | Other |
| UPMC-HNC | 308 | 81 | 7 | 2061102 | no evidence of disease (NED): **197**<br>non-NED: **111** | 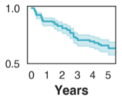 | recurred: **31**<br>not recurred: **253** | N/A | HPV infected: **158**<br>not HPV infected: **150** |



### Supplemental Methods

##### CODEX data collection

All samples are prepared, stained, and acquired following CODEX User Manual Rev C (https://www.akoyabio.com).

**Coverslip preparation:** Coverslips are coated with 0.1% poly-L-lysine solution to enhance adherence of tissue sections prior to mounting. The prepared coverslips are washed and stored according to the guidelines in the CODEX User Manual.

**Tissue sectioning:** formaldehyde-fixed paraffin-embedded (FFPE) samples are sectioned at a thickness of 3-5 μm on the poly-L-lysine coated glass coverslips.

**Antibody conjugation:** Custom conjugated antibodies are prepared using the CODEX Conjugation Kit, which includes the following steps: (1) the antibody is partially reduced to expose thiol ends of the antibody heavy chains; (2) the reduced antibody is conjugated with a CODEX barcode; (3) the conjugated antibody is purified; (4) Antibody Storage Solution is added for antibody stabilization for long term storage. Post-conjugated antibodies are validated by SDS-polyacrylamide gel electrophoresis (SDS-PAGE) and quality control (QC) tissue testing, where immunofluorescence images are stained and acquired following standard CODEX protocols, then evaluated by immunologists.

**Staining:** CODEX multiplexed immunofluorescence imaging was performed on FFPE patient biopsies using the Akoya Biosciences PhenoCycler platform (also known as CODEX). 5 μm thick sections were mounted onto poly-L-lysine-treated glass coverslips as tumor microarrays. Samples were pre-treated by heating on a 55 ℃ hot plate for 25 minutes and cooled for 5 minutes. Each coverslip was hydrated using an ethanol series: two washes in HistoChoice Clearing Agent, two in 100% ethanol, one wash each in 90%, 70%, 50%, and 30% ethanol solutions, and two washes in deionized water (ddH2O). Next, antigen retrieval was performed by immersing coverslips in Tris-EDTA pH 9.0 and incubating them in a pressure cooker for 20 minutes on the High setting, followed by 7 minutes to cool. Coverslips were washed twice for two minutes each in ddH2O, then washed in Hydration Buffer (Akoya Biosciences) twice for two minutes each. Next, coverslips were equilibrated in Staining Buffer (Akoya Biosciences) for 30 minutes. The conjugated antibody cocktail solution in Staining Buffer was added to coverslips in a humidity chamber and incubated for 3 hours at room temperature or 16 hours at 4 ℃. After incubation, the sample coverslips are washed and fixed following the CODEX User Manual.

**Data acquisition:** Sample coverslips are mounted on a microscope stage. Images are acquired using a Keyence microscope that is configured to the PhenoCycler Instrument at a 20X objective. All of the sample collections were approved by institutional review boards.

To correct for possible autofluorescence, “blank” images were acquired in each microscope channel during the first cycle of CODEX and during the last. For these images, no fluorophores were added to the tissue. These images were used for background subtraction. Typically, autofluorescence will decrease over the course of a CODEX experiment (due to repeated exposures). Thus, to correct each cycle, our method determines the extent of subtraction needed by interpolating between the first and last “blank” images.
